# Potent neutralization of clinical isolates of SARS-CoV-2 D614 and G614 variants by a monomeric, sub-nanomolar affinity Nanobody

**DOI:** 10.1101/2020.06.09.137935

**Authors:** Guillermo Valenzuela Nieto, Ronald Jara, Daniel Watterson, Naphak Modhiran, Alberto A Amarilla, Johanna Himelreichs, Alexander A. Khromykh, Constanza Salinas, Teresa Pinto, Yorka Cheuquemilla, Yago Margolles, Natalia López González del Rey, Zaray Miranda-Chacon, Alexei Cuevas, Anne Berking, Camila Deride, Sebastián González-Moraga, Héctor Mancilla, Daniel Maturana, Andreas Langer, Juan Pablo Toledo, Ananda Müller, Benjamín Uberti, Paola Krall, Pamela Ehrenfeld, Javier Blesa, Pedro Chana-Cuevas, German Rehren, David Schwefel, Luis Ángel Fernandez, Alejandro Rojas-Fernandez

## Abstract

Despite unprecedented global efforts to rapidly develop SARS-CoV-2 treatments, in order to reduce the burden placed on health systems, the situation remains critical. Effective diagnosis, treatment, and prophylactic measures are urgently required to meet global demand: recombinant antibodies fulfill these requirements and have marked clinical potential. Here, we describe the fast-tracked development of an alpaca Nanobody specific for the receptor-binding-domain (RBD) of the SARS-CoV-2 Spike protein with therapeutic potential applicability. We present a rapid method for nanobody isolation that includes an optimized immunization regimen coupled with VHH library *E. coli* surface display, which allows single-step selection of high-affinity nanobodies using a simple density gradient centrifugation of the bacterial library. The selected single and monomeric Nanobody, W25, binds to the SARS-CoV-2 S RBD with sub-nanomolar affinity and efficiently competes with ACE-2 receptor binding. Furthermore, W25 potently neutralizes SARS-CoV-2 wild type and the D614G variant with IC50 values in the nanomolar range, demonstrating its potential as antiviral agent.

## Introduction

Severe clinical courses of pandemic coronavirus disease 2019 (COVID-19), the illness caused by SARS-CoV-2 infection, involve pneumonia and multiple organ dysfunction, and constitute an unprecedented threat to health and economy worldwide (1-4). Currently, there are no vaccines or drugs to effectively contain the pandemic. In order to avoid the collapse of healthcare systems, non-pharmaceutical public health measures such as social distancing, border closures, and lockdowns have been enforced globally (5, 6). Genetic studies determined that the pathogen responsible for this outbreak belongs to the Coronaviridae family, genus *Beta-coronavirus*, sub-genus *sarbecovirus (7)*. It has high sequence homology with the bat coronavirus RaTG13, indicating that the novel virus may have originated in bats and subsequently jumped to humans, probably via a yet unidentified intermediate animal host (8).

The positive sense SARS-CoV-2 RNA genome contains 29903 nucleotides, including 12 open reading frames (ORFs) coding for the replicase ORF1ab polyproteins, Spike, Envelope, Membrane and Nucleocapsid structural proteins, and several accessory proteins (9, 10). The Spike protein on the virion surface is responsible for attachment to, and invasion of host cells (11). Spike is a highly glycosylated trimeric class I fusion protein and contains two subunits, S1 and S2 (12). Similar to SARS-CoV, Angiotensin-converting-enzyme 2 (ACE2) appears to be the molecular entryway to the host, since SARS-CoV-2 S binds to this receptor (12-15). Moreover, the presence of the ACE2 receptor has been confirmed in a variety of human tissues that are related to clinical manifestations of COVID-19 (16-19).

CryoEM studies showed that SARS-CoV-2 Spike exhibits a metastable pre-fusion conformation, where the RBD within S1 performs hinge-like movements between “down”- and “up”-positions relative to the remainder of the S protein. Only in the “up”-position, RBD residues responsible for binding to the ACE2 receptor on the host cell surface are exposed. After attachment, proteolytic processing and S1 shedding, the S2 subunit undergoes substantial conformational re-arrangements to a stable post-fusion conformation, concomitant with membrane fusion and invasion of the host cell (20, 21). Host TMPRRS2 serine proteases seem to be responsible for this proteolytic priming, attacking a furin-like cleavage site situated in between the S1 and S2 subunits of the Spike protein (22, 23).

Recent studies highlighted the emergence of the Spike protein variant D614G, which has become the dominant SARS-CoV-2 pandemic strain (24). This variant seems to replicate better in cell culture, but it is disputed if the mutation results in increased viral load or infectivity in humans (25, 26). Interestingly, the S protein residue 614 is located in the interface between adjacent S protomers, and it has been hypothesized that amino acid exchange to glycine stabilizes the trimeric Spike protein architecture (27) Accordingly, the D614G variant exhibits less S1 subunit shedding and improved Spike protein incorporation into virions (28). However, the mutation does not influence receptor binding or antibody neutralization, and seems not to be associated with worse clinical outcome (29, 30)

Altogether, the central role of the Spike glycoprotein in the virus lifecycle highlights the importance of this protein as a target for the development of therapies such as neutralizing antibodies and vaccines (31-35). In this sense, isolation of specific Spike protein antibodies can be instrumental in the development of effective diagnostic and therapeutic tools (36-40).

Some naturally occurring antibodies that lack light chains (heavy-chain only antibody, HCAb) are known as single-domain antibodies. They are derivates of IgG and occur in the entire camelidae family (41). The camelidae family consists of camels, dromedaries, llamas, vicuñas, guanaco, and alpacas (42). The antigen-binding fragment of an HCAb contains a single variable heavy homodimer (VHH) domain consisting of 3 hypervariable regions. An isolated VHH domain is also referred to as a Nanobody. Nanobodies can be used as therapeutic bullets for instance, against tumors, pathogens, and chronic diseases (43-45). Nanobodies are in general more stable than conventional antibodies and, unlike classical antibodies, can be efficiently produced in prokaryotic expression systems. In fact, several milligrams can be produced from one liter of culture (43, 46-49), offering a means to rapidly and economically produce therapeutic biologics at large scale. Nanobodies derived from camelid HCAbs are obtained after immunization with the target protein plus adjuvant. Our platform has developed an improved procedure to produce Nanobodies using alpacas as the donor species. To obtain the genetic sequences of target-specific Nanobodies produced after immunization, peripheral blood B-lymphocytes are isolated to obtain total RNA, followed by cDNA preparation to finally amplify the Nanobody region (50). The cDNA fragment encoding the Nanobody is as short as 360 nt, and up to ∼ 3×10^6^ single clones can be obtained from 120 mL blood in a safe and harmless procedure for our alpacas. Afterward, a bacterial display system is applied to rapidly clone the full single Nanobodies. Here, we make use of the bacterial display system to develop a fast, inexpensive and simple density gradient method for Nanobody selection, by which we identified a Nanobody against SARS-CoV-2 with strong binding and neutralizing activity.

## Results

### Immunization and density gradient method for Nanobody selection

We developed a method to fast-track Nanobody selection from alpacas and use the approach to obtain a Nanobody against the Spike protein of SARS-CoV-2 in only few weeks. We used lyophilized Spike protein produced in a baculovirus expression system as antigen. Prior to immunization, antigen integrity was tested by SDS-Page and Coomassie staining (Figure 1a). Then, an alpaca (Figure 1b) was immunized twice over 14 days with 100 μg of the full Spike protein and adjuvant. The animal′s health was monitored throughout the study period by clinical examination, hematology analysis, and serum biochemistry. The immune response of the alpaca′s serum before immunization revealed a fortunate basal cross-reaction against the Spike protein, and after the second immunization, we observed a significant increase of antigen specific IgG antibodies in the alpaca′s serum. This analysis was done in a rapid qualitative manner by Dot blot analysis, immobilizing the epitope to a nitrocellulose membrane and using alpaca serum as a source of primary antibodies (Figure 1c). We also detected an ∼ 5-fold increment of IgG antibodies against Spike protein in the post-immunization serum by ELISA (Figure 1d). Our method for Nanobody isolation is based on the bacterial display system, a strategy that takes advantage of the high transformation efficiency of *E. coli* and avoids the need for bacteriophage infections or shuttling into yeast cells for surface display of the Nanobodies (51). Importantly, this Nanobody display system can drive the specific adhesion of *E. coli* bacteria to abiotic and cellular surfaces with the cognate antigen (52, 53). Thus, we constructed a bacterial display library with a complexity of 2.3 ×10^6^ independent clones by electroporation of *E. coli* DH10B-T1R strain (53-55) (see Materials and Methods).

**Figure 1.**
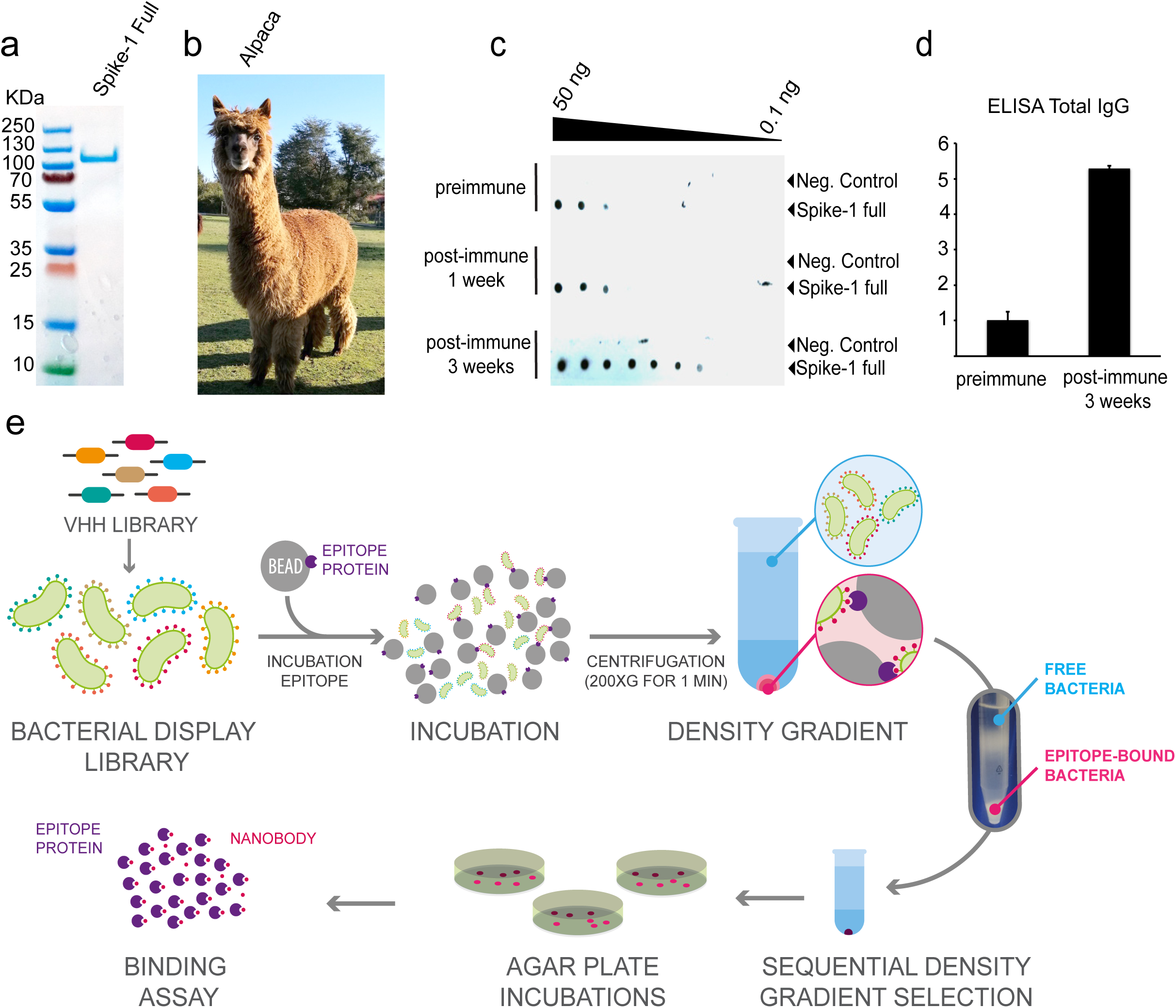
Immunization of the Spike of SARS-CoV-2 and a simple density gradient method for the selection of Nanobodies. **A)** SDS-Page to ensure protein integrity of full-length Spike of SARS-CoV-2 before immunization. **B)** Adult alpaca immunized with Spike. **C)** Evaluation of the alpaca′s immune response by Dot blot. Image shows the reaction to decreasing amounts of Spike-1 and bovine serum albumin (negative control) using a pre-immunization control, and after one immunization (1 week), or two immunizations (3 weeks) with full-length SARS-CoV-2 Spike, using alpaca serums as a primary antibody source followed by an anti-camelid IgG-HRP secondary antibody. **D)** ELISA assay before and after the second immunization (3 weeks). **E)** Schematic representation of novel protocol for isolation of Nanobodies using density gradient separation. The bacterial display library expressing the Nanobodies on the surface of bacteria is briefly incubated with conventional sepharose beads coated with the epitope of interest. Directly after the mixture is deposited on a Ficoll gradient conic tube and centrifuged at 200 g for 1 min, the beads drive through the gradient to the bottom of the tube with the bacteria expressing specific Nanobodies, while unbound bacteria remain on the surface of the gradient. The beads are then resuspended, and bacterial clones are isolated for biochemical binding confirmation.

We applied a novel procedure for the selection of Nanobodies based on a simple Ficoll density gradient, an inexpensive reagent available all around the world used for blood fragmentation. We were inspired by the main observation that red cells accumulate at the bottom of the Ficoll density gradient, while PBMCs stay in the upper fraction. Using conventional NHS-activated sepharose beads in a Ficoll gradient, we found that the density of the beads was sufficient to precipitate to the bottom of a 15 mL tube. Furthermore, the same assay was performed with free bacteria, and as expected, the bacteria remained on top of the gradient. In the bacterial display system, a single Nanobody clone is expressed by each bacterium of the library (51). *E. coli* bacteria express intimin-Nanobody protein fusions that anchor in the outer membrane upon IPTG-induction and expose the functional Nanobody to the extracellular milieu for antigen recognition. These intimin-Nanobody fusions also contain a common epitope (myc-tag) at the C-terminus for immunodetection (54). Therefore, the bacteria expressing specific Nanobodies on their surface would bind NHS-beads coated with the protein of interest (i.e. SARS-CoV-2 S) and migrate all the way through to the bottom of the Ficoll density gradient, leaving unbound bacteria in the upper fraction (Figure 1e). Indeed, we demonstrated that specific Nanobodies from a bacterial display library are rapidly selected with our protocol, using common, inexpensive reagents and a conventional centrifuge. A full description is provided in the Materials and Methods section.

### Nanobody isolation and detection of Spike of SARS-CoV2 by IF and ELISA assays

We optimized conditions to extract the intimin-Nanobody fusions from the bacterial outer membrane and used these protein extracts directly for assaying the binding to the Spike protein applying two different methods, dot blot analysis and high-content microscopy. After Nanobody selection using our simple Ficoll-based density gradient protocol, we obtained ∼ 1000 colonies on LB-agar plates from the sepharose-antigen coated fraction. 100 colonies were analyzed of which 30 showed a high affinity for Spike. The bacteria were inoculated in liquid LB media and the expression of intimin-Nanobodies of the single clones was induced. Bacterial pellets were lysed under optimized conditions and the extract was used as a source of Nanobodies for the second binding screening. For dot blot analysis, a negative control of an unrelated protein was applied adjacent to the same amount of full-length Spike protein on the nitrocellulose strips. Single Dot blot strip tests were incubated with the bacterial extracts containing Nanobodies. Sequential incubation with mouse anti-myc antibody and an anti-mouse HRP-conjugate unveiled nanobodies binding to Spike. Of the first 100 clones, we focused on two Nanobodies that displayed a strong signal for full-length Spike in the Dot blot analysis (W23 and W25, Figure 2a). Additionally, we used high content microscopy as a second confirmation method. For this purpose, a single 10 cm-plate was transfected with a Spike-GFP vector for 24h and further the cells were seeded onto 96 well-plates. After 24 hours, the cells were fixed, permeabilized, and individual extracts of our 100 selected bacterial display clones were added as a source of Nanobodies. A mouse anti-myc antibody and an anti-mouse Alexa647 secondary antibody was used for immunofluorescence detection. HeLa cells had a transfection efficiency of ∼20%. In this case, a low transfection rate is desired, because it allows us to evaluate unspecific binding to un-transfected cells in the same image. Consistent with the Dot blot analysis, the W23 and W25 Nanobodies bound to the Spike-GFP expressed in human cells (Figure 2b). We observed co-localization of W23 and W25 to Spike-GFP, while no co-localization was observed with negative control extracts (Figure 2b). We also observed that W25 does not bind to the nucleoprotein of SARS-CoV-2 tagged to a GFP protein (Supplemental Figure 1b) confirming that W25 binds specifically to Spike and not to GFP. The selected clones were sequenced and the alignment of the amino acid sequences showed a high similarity between the two Nanobodies, CDR3 was identical, suggesting that the two Nanobodies are members of the same family and were most likely generated from a common origin during the Alpaca′s immunoreaction against the Spike protein (Figure 2c). In conclusion, we developed a method that rapidly performs secondary screening selection of Nanobodies, using bacterial extracts from selected clones of the bacterial display library, using either conventional biochemical methods such as dot blot analysis, or high content microscopy immunofluorescence-based assays.

**Figure 2.**
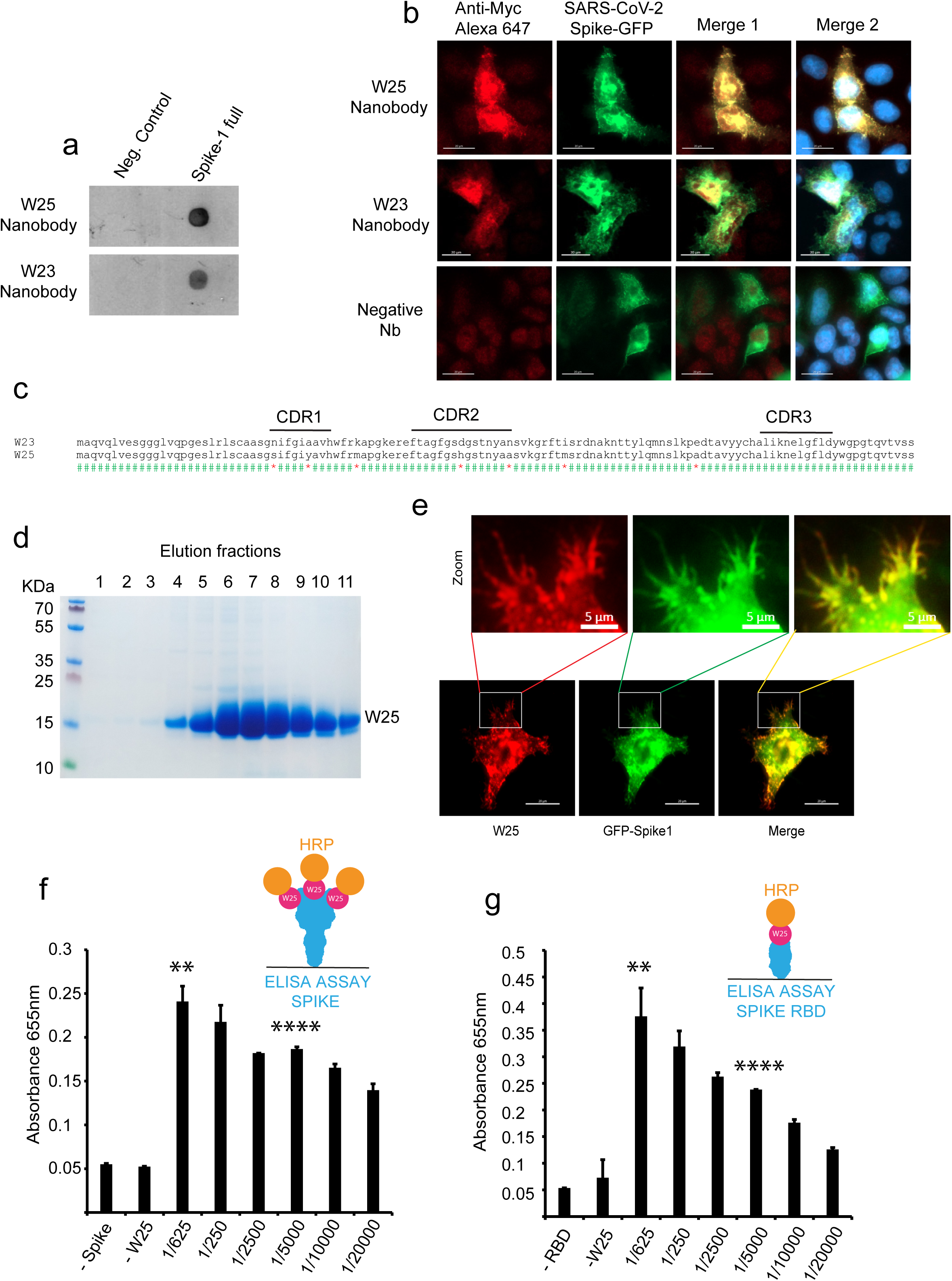
Dual biochemical and microscopy-based selection of Nanobodies. **A)** Dot blot immunodetection of full-length SARS-CoV-2 Spike using direct total protein extracts of clones W25 and W23 as the primary antibody. Mouse anti-Myc (1:3000) followed by anti-mouse-HRP were used for detection. Protein extract from *E. coli* (BL21 strain) was used as a negative control. **B)** Immunodetection of Spike-GFP transiently transfected in HeLa cells using total protein extract selected clones as the primary antibody, followed by mouse anti-Myc (1:3000) and anti-mouse-Alexa 647. The image shows two positive clones (W25 and W23), and an example of a negative Nanobody. **C)** Sequence alignment of aminoacidic sequence of W25 and W23. CDR sequences are marked with a black line. **D)** Purification from periplasm of bacteria, elution fraction of a single liter of bacterial culture. **E)** Immunodetection as in B, using purified protein. **F)** ELISA assay of full-length Spike of SARS-CoV-2 using conjugated W25-HRP Nanobody **G)** ELISA assay of RBD of Spike using W25-HRP conjugate Nanobody. Statistic t-test using software R 1.2.1335, ** P ≤0.01; **** P ≤0.0001 compare to -Spike control

W25 was subsequently cloned into the pHen6 vector for periplasmic bacterial expression, and large amounts of recombinant W25 Nanobody were obtained (Figure 2d). The purified W25 Nanobody was used for immunofluorescence of HeLa cells transiently transfected with an S-GFP vector and specifically recognized the Spike-GFP in a sensitive manner (Figure 2e), given that we observed co-localization of Spike-GFP and W25 to membrane protrusions in HeLa cells (Figure 2e). Thus, our experiment suggests the W25 Nanobody is applicable as a measure for the direct diagnosis of infected cells by immunofluorescence. Furthermore, we covalently labelled W25 to Horseradish Peroxidase (HRP) and performed direct ELISA assays using immobilized full-length Spike protein (Figure 2f) or RBD only (Figure 2g). In both cases, W25 was able to recognize in an efficiently and sensitivemanner the viral proteins immobilized on the ELISA plate. Due to the high levels of expression and effective conjugation to HRP, the Nanobody W25 will be a convenient tool for the development of diagnostic approaches based on direct antigen detection. Furthermore, the efficient binding to RBD might indicate SARS-CoV-2 neutralization activity.

### Sub-nanomolar affinity recognition of RBD and efficient competition for ACE2 by the W25 Nanobody

We further characterized the direct interaction between the Spike RBD and the W25 nanobody by pulldown assays, employing recombinant RBD protein or BSA as a control. Proteins were covalently immobilized on NHS-sepharose beads, and the binding of W25 to RBD and control beads was tested. We showed that the selected W25 Nanobody binds to RBD (Figure 3a) confirming previous ELISA results. To further study the interaction of W25 with S RBD, we first used thermal shift assays. We demonstrated integrity of the individual RBD and Nanobody preparations using a label-free Tycho measurement, yielding inflection temperatures of protein unfolding (T_i_) of 52.4°C and 57.9°C, proving high thermal stability. Mixing the RBD domain with the nanobody, followed by the same measurement, led to a shift to higher unfolding inflection temperature of ∼14°C, strongly indicative of tight interaction between the RBD and Nanobody (Figure 3b). To quantify the interaction affinity between the RBD domain and W25, Microscale Thermophoresis (MST) was employed. Fitting of the experimental MST fluorescence traces resulted in a binding affinity of W25 to the S RBD of 295 pM ± 84 pM (Figure 3c). We furthermore tested whether W25 and the soluble domain of the ACE2 receptor would compete for binding to Spike RBD. We labelled W25 with a florescent dye and generated and stable complex with RBD at a very low concentration of 1 nM W25 and 2 nM RBD. Increasing amounts of ACE2 were added, and dissociation of W25 was determined by thermophoresis. We observed an effective competition between W25 and ACE2 for RBD, where the ACE2 concentration to reach half-maximal W25-dissociation (EC50) was determined to be 33 nM ± 9 nM (Figure 3d, e). These results indicate that the binding affinity of the Nanobody W25 for RBD is considerably higher than that of ACE2, and suggest a direct competition for RBD interface residues between W25 and ACE2.

**Figure 3.**
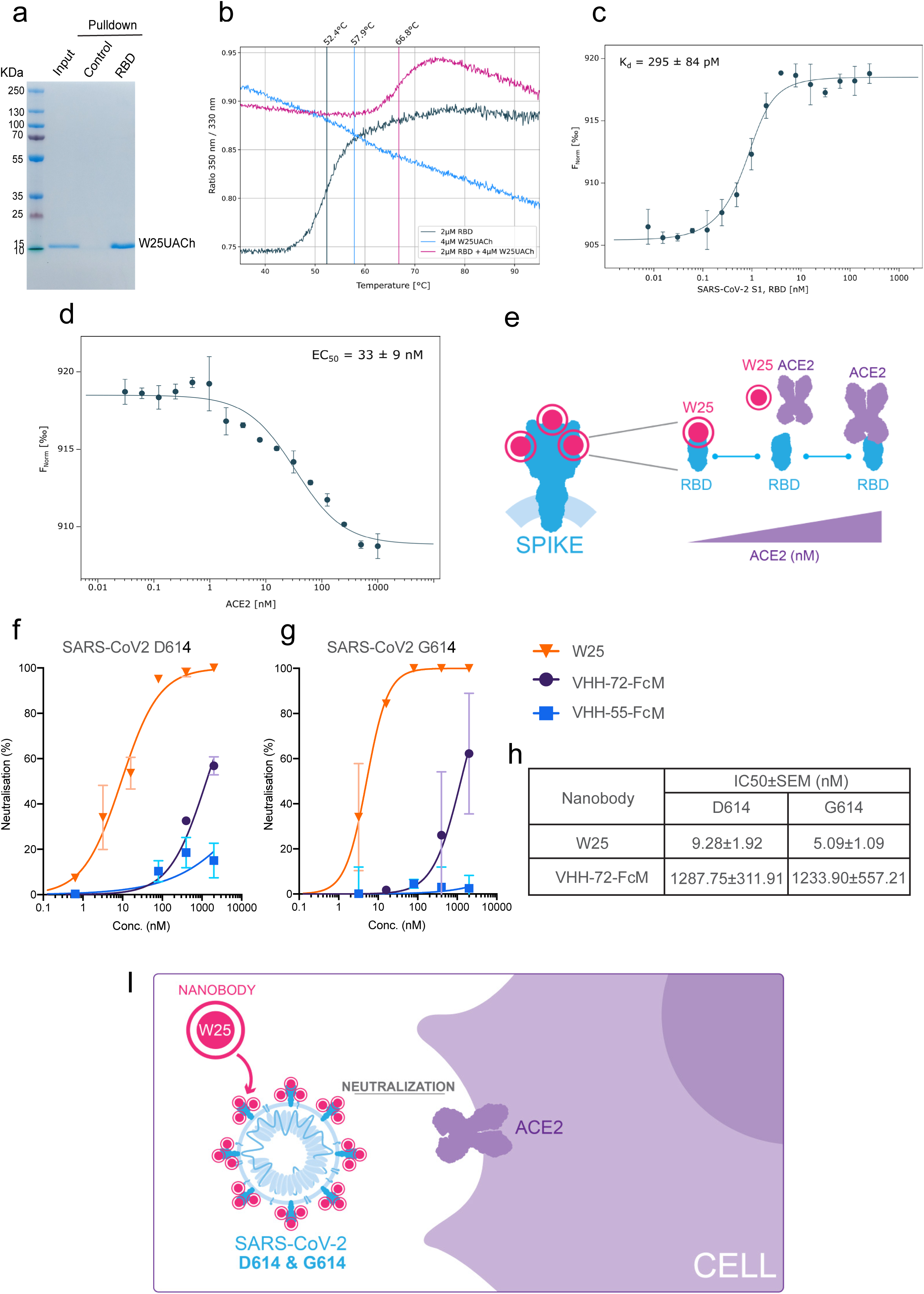
Binding characterization and Neutralization of SARS-CoV-2 by the Nanobody W25. **A)** Pulldown of the W25 Nanobody. A recombinant Spike RBD domain of the SARS-CoV-2 Spike protein or control BSA protein were covalently bound to NHS-sepharose beads. Further, the W25 Nanobody was incubated with control and Spike RBD beads, washed, and further eluted in LSD lysis buffer (Invitrogen). Detection was performed by Coomassie staining. **B)** Unfolding profiles of 2 μM SARS-CoV-2 S1, Spike RBD in the absence (black) and presence (red) of 2 μM W25, measured with Tycho NT.6. Binding of W25 to Spike RBD leads to strong stabilization and shifts the inflection unfolding temperature (T_i_) from 52.1°C to 66.3°C. **C)** MST binding curve for the titration of 1 nM fluorescently labeled W25 into a 16-point serial dilution of SARS-CoV-2 S1, Spike RBD (250 nM – 7.6 pM). W25 binds Spike RBD with sub-nanomolar affinity (K_d_ = 295 ± 84 pM). Error bars show the SD calculated from experiments performed in triplicate. **D)** MST competitive curve for 2 nM of fluorescently labeled W25UACh incubated with 4 nM SARS-CoV-2 RBD, titrated with a 16-point dilution series of hACE2 (1 μM – 30.5 pM). W25UACh is displaced by hACE2 with nanomolar concentration (EC50 = 33 ± 9 nM). Error bar show the SD calculated from triplicate experiments. **E)** Diagram of W25 and ACE2 competition for RBD of Spike of SARS-CoV-2. **F)** Neutralization efficiency of the Nanobody W25 was compare to the previous reported Nanobodies VHH-72-Fc monomeric and VHH-55-Fc monomeric. **G)** Neutralization assays with SARS-CoV-2 wt “D614 variant”. **H)** Comparative EC50 values for neutralization by W25 and VHH-72 FcM against D614 and G614 virus variants. **I)** Diagram of W25 neutralization of SARS-CoV-2

### W25 as single domain strongly neutralizes SARS-CoV2 viruses

Next, we investigated the neutralization ability of W25 against two clinical isolates of SARS-CoV2 infections, Spike protein variants D614 and G614, in Vero E6 cells. W25 had demonstrated strong neutralizing activity with IC50 values of 9.82 nM ± 1.92 nM and 5.09 nM ±1.09 nM for the D614 and G614 SARS-CoV-2 variant, respectively. The SARS-CoV-2 and MERS-specific Nanobodies, VHH-72 and VHH-55 (39), produced as monomeric Fc fusions, were included in the measurements for comparison. VHH-72 exhibited a moderate degree of neutralization with an IC50 of 1287.75 nM ± 311.91 nM and 1233.90 nM ± 557.21 nM for S protein D614 and G614 variants, respectively. As expected, no inhibition was observed for the MERS-specific VHH-55. Thus, W25 represents a potent neutralizing monomeric Nanobody against SARS-CoV-2, with improved activity relative to other described HCAbs, and is able to inhibit the predominant circulating virus strain containing the Spike protein D614G mutation (Figure 3f-i).

## Discussion

Currently, selection of Nanobodies from bacterial display is performed by affinity purification based on magnetic beads binding the labelled antigen or by FACS Sorter or selection on cells transfected with the antigen (55). Here, we describe a novel simple method for the selection of Nanobodies from *E. coli* bacterial display libraries. Our method requires only conventional laboratory instruments and an inexpensive Ficoll gradient for the selection of Nanobodies from a highly complex and diverse bacterial library. The success rate depends on the immune response of the alpaca to the epitope and by prior undetermined exposure to cross-reactive camelid coronaviruses (56-58). Nevertheless, here we have shown a successful example based on an immunization program of two weeks duration and 100 analyzed clones resulting in the selection of 30 Nanobodies with high affinity for the antigen; thus, our method can accelerate the identification of Nanobodies, enabling, for example, the generation of diagnostic and potentially therapeutic measures against Covid-19 and other infectious and emergent viruses.

The medical, social, and economic consequences of COVID-19 impact the lives of everyone on the planet and while vaccines and effective therapies are not available, sensitive diagnostic tools are urgently needed. Currently, convalescent plasma transfusion has been applied as an emergency treatment for COVID-19, aiming to enhance the immune response of the patients (59, 60). In the late phase of disease progression, an inflammatory cytokine storm has been reported. Antibody administration at this stage by convalescent plasma transfusion is under discussion, especially for critically ill patients (61). Also additional efforts have been done for the generation and clinical implementation of hyperimmune equine serum therapy approaches (62). Nanobodies might become an alternative for immunotherapy to replace or complement convalescent plasma transfusion and equine hyperimmune serum. Nanobodies can be rapidly and cost-effectively produced in an active form in prokaryotic systems, and in addition the lack of an Fc domain reduces the possibility of undesired immunostimulatory activity on the host.

Here, we characterize the affinity properties of novel Nanobodies against the Spike protein of SARS-CoV-2. We demonstrate that the Nanobody W25 can be suitable as a diagnostic reagent and show that it is able to sensitively detect Spike by immunofluorescence in human cells as well as for direct Spike determination by ELISA Assays. The yield of production of W25 is high (∼ 60mg/L in bacterial culture) as shown in Figure 2d and therefore inexpensive. Using fast thermal denaturation, we show that W25 interacts with the Spike RBD. We further determined by Thermophoresis that W25 has an affinity for the RBD domain of Spike of Kd ∼0.3 nM. To date, four research groups have reported Nanobodies and synthetic antibodies against Spike RBD: VHH 72 with an Kd of 39nM (39); Ty1 with an Kd of 8 ± 1.5 nM at normal salt conditions (63); a large set of eighteen humanized single-domain antibodies, with the highest affinity clone n3021 demonstrating a Kd of 0.63 ±0.01 nM (36); the H11-H4 Kd of 5nM (64); and the recently synthetic monomeric mNb6 with an Kd of 0.56 nM (65), the latter also shows enhanced affinity by trimerization as expected by an additive increment of the avidity for the trimeric Spike protein. Thus, to date the neutralizing monomeric W25 Nanobody reported here shows the highest affinity for the RBD Spike protein of SARS-CoV-2. We have also demonstrated efficient neutralization mediated by W25 against clinical isolates of live viruses. An early isolate, similar to the original virus found in Wuhan, was neutralized by the W25 monomeric nanobody with an IC50 ∼9.28 nM. Interestingly a slightly enhanced neutralization effect was observed against the currently dominant SARS-CoV-2 variant G614, with an IC50 of ∼5.09 nM. As residue 614 is outside the RBD, this finding may suggest differential RBD exposure between the two strains, which could underpin the observed selection for the G614 over time.

Animal testing for *in vivo* safety and efficacy in animals would be required to ascertain the therapeutic potential of W25. Our aim is to provide a stable, and scalable production of nanobody for the generation of a neutralizing inhaler able to block the viral replication directly in the upper airway in the early stages of the Covid-19 development. Nanobodies have been used previously with the same approach to treat syncytial virus infection (RSV). Successful preclinical and clinical trials have been performed indicating that 6mg/kg has been a safe and efficient dose for RSV (66-70). In this case, a monomeric Nanobody called Nb017 with a Kd of ∼ 17.88 nM was trimerized to a drug called ALX-0171 increasing the binding affinity to RSV to a Kd of ∼ 0.113 nM. Since W25 already shows strong neutralizing activity against two important clinical virus isolates as monomer, such approaches could directly enhance efficacy and delivery. Interestingly, a recent study reported that the more prevalent virus variant G614 is associated with higher levels of viral nucleic acid in the upper respiratory tract in human patients, potentially making it even more accessible for inhaler delivery (24).

## Material and Methods

### Immunization and VHH library construction

The alpaca immunization process followed the guidelines established by the Bioethics Committee of the Austral University of Chile (certifications 338/2019 and 388/2020). One day before immunization, 5 mL of blood was collected for pre-immune serum tests. For immunization (day 1), 100 μg of full-length Spike protein of SARS-CoV-2 (SINOBiological) was used. The cold lyophilized protein was dissolved in 2 mL of adjuvant (Veterinary Vaccine Adjuvant, GERBU FAMA) diluted 1:1 in sterile water and injected subcutaneously into a male alpaca (*Vicugna pacos*). A total volume of 4 mL was injected into four different locations in the alpaca. A 5 mL blood sample was collected seven days after the first immunization. On day 14, the alpaca was immunized again with 100 μg Spike, and on day 15 a sample of 120 mL blood was collected from the jugular vein in tubes containing 3.8% sodium citrate as an anti-coagulant. The uncoagulated blood sample was mixed with the same volume of HBSS medium without calcium (Gibco), divided into aliquots of 10 mL and each aliquot was added on top of a 5 mL of Ficoll-Paque Premium (GE Healthcare) in 15 mL sterile Falcon tubes. After centrifugation (1,200 rpm, 80 min, RT), the PBMC fraction was recovered from the interphase, washed twice in PBS by centrifugation (3,500 rpm, 10 min), resuspended in 4 mL of sterile PBS (Gibco). RNA extraction and cDNA production were performed using the commercial RNeasy Mini Kit (Qiagen) and QuantiTect Reverse Transcription Kit (Qiagen) respectively according to the manufacturer’s instructions. Approximately 2 μL of each cDNA synthesis procedure were used as templates in 50 μL PCR reactions with oligonucleotides CALL001 (5′-GTC CTG GCT CTC TTC TAC AAG G-3′) and CALL002 (5′-GGTACGTGCTGTTGAACTGTTCC-3′) (71). The amplified fragments of ∼0.6 kb, corresponding to VHH-CH2 domains, and ∼0.9 kb, corresponding to conventional VH-CH1-CH2 domains, were separated in 1.2% (w/v) low melting agarose gels and the ∼0.6 kb band was purified (QIAEX II Gel Extraction kit, Qiagen). This fragment was used as a template in a second PCR reaction with oligonucleotides VHH-Sfi2 (5′-GTC CTC GCA ACT GCG GCC CAG CCGGCC ATG GCT CAG GTG CAG CTG GTG GA-3’) and VHH-Not2 (5′-GGA CTA GTG CGG CCG CTG AGG AGA CGG TGA CCT GGG T-3′) to finally obtain the amplified fragments of ∼0.4 kb, corresponding to VHH domains. The amplified VHH fragments were digested with SfiI and NotI restriction enzymes (Thermo Scientific) and ligated into the same sites of the purified vector pNeae2 (55). Ligations were electroporated in *E. coli* DH10B-T1 R cells obtaining a library size of ∼3 × 10^6^ individual clones, as determined by plating on LB-chloramphenicol agar plates with 2% w/v glucose incubated at 30°C. Less than 0.7% of re-ligated vectors were estimated from a control ligation performed in parallel without the DNA insert. Transformed bacteria were scraped from plates and stored at -80°C in LB broth with 30% glycerol.

### Coupling epitopes to beads

1 mL of NHS-activated sepharose 4 Fast Flow beads were washed with 2 mL of cold 1 mM HCI immediately before use, then washed 5 times with cold sterile PBS. 200 μg of purified protein in PBS 1X was added to the beads and incubated with rotation overnight. Non-reacted groups in the medium were blocked by adding ethanolamine to a final concentration of 0.5 M. Beads were washed 5 times with PBS and stored at 4°C.

### ELISA Assays

*For total IgG detection of alpacas serum*, 10 ng of SARS-CoV-2 Spike protein diluted in PBS 1X pH 7.4 were bound in each well of the ELISA plate, incubated at 37°C for 1h. Followed by a 3 × 5 min wash with PBS-T (PBS, 0.1% Tween20). Serial dilutions (1:5000 in PBS) of pre and post immunization serum were incubated at 37°C for 1h. Followed by a 3 × 5 minutes wash with PBS-T. Secondary detection was done with peroxidase-labeled anti-Llama IgG (1:5000 in PBS-T), followed by a 3 × 5 min wash with PBS-T and visualized using 1-step Ultra TMB-ELISA (THERMO), 5 min at 37°C. Signal was measured at 650nm in a microplate reader after 10 min incubation. *For direct detection of Spike and RBD by covalent conjugate* W25, 5mg of W25uach nanobody was conjugated with horseradish peroxidase (HRP) using Thermo Scientific Pierce Maleimide Activated Horseradish Peroxidase Kit, according manufacturer instructions in order to covalently attach HRP to sulfhydryl groups from nanobody cysteines. 96-well ELISA plates were coated with 100ng of full-length Spike protein or RBD only diluted in PBS 1X pH 7.4, incubated 1 hr at 37°C, followed by 3 × 5 min wash with PBS-T. Serial dilutions of W25uach-HRP in PBS 1X were incubated at 37°C for 1h in humid chamber. Followed by a 3 × 5 minutes wash with PBS-T and visualized using 100μL of 1-step Ultra TMB-ELISA (THERMO), 5 min at 37°C. Signal was measured at 650nm in a microplate reader.

### Density gradient separation

1 mL of glycerol stock from the library was inoculated in a flask containing 20 mL of LB medium with 25 μg/mL of chloramphenicol and 2% glucose. The flask was incubated (pre-inoculum) overnight at 37°C with 200 rpm agitation. The same procedure was repeated with control bacteria that were transformed with a kanamycin resistant plasmid (negative control). The pre-inoculum was pelleted and resuspended in LB medium with 25 μg/mL chloramphenicol and then diluted to OD600=0.02 in 100 mL of fresh LB medium with 25 μg/mL chloramphenicol without glucose, incubated at 37°C with 200 rpm agitation until it reached 0.45 - 0.6 OD600. IPTG was added to a final concentration of 50 μM to induce protein expression for 3 hours at 30°C and 200 rpm. OD600 absorbance of library and control bacteria cultures was measured. 50 mL of both cultures was washed three times with 10 mL of filtered PBS. Centrifugation was always performed at 3,000 g for 5 min. Library culture (chloramphenicol resistant) and negative control (kanamycin resistant) were resuspended in a final volume of 10 mL PBS. 2 mL of library culture and 2 mL of negative control cultures were mixed (if OD600nm was different, the volume of the control bacteria was adjusted based on OD in order to add an equal amount of bacteria) and incubated with 300 μL of NHS beads coupled to epitope protein on a 15 mL conical tube on a rocking platform for 30 min at room temperature. The mixture was slowly added on the top of 6 mL of Ficoll (Ficoll-PaqueTM PLUS GE Healthcare) in a 15 mL conical tube, centrifuged at 200 x g for 1 min. The unbound fraction was discarded (upper fractions), leaving a visible pellet of beads that was resuspended in 4 mL PBS and rotated for 5 min at room temperature. This step was repeated six times. Finally, 1 mL of LB medium was added and incubated for 5 min at room temperature, then 50 μL were plated on LB agar plates with 50 μg/mL kanamycin and 2% glucose, 50 μL were plated on LB agar plates with 25 μg/mL chloramphenicol and 2% glucose and the rest in at least two LB chloramphenicol/glucose agar plates, incubated at 37°C overnight (>20hrs recommended). The colony number of the first two plates were counted as a measurement of specific enrichment of Nanobodies expressing bacteria from the library.

### Expression, sub-cloning and protein purification

The selected VHH cDNA fragments were digested with SfiI and NotI restriction enzymes (Thermo Scientific) and ligated into the same sites of the purified vector pHen6 (72). For periplasmic expression the *E. coli* wk6 strain was used. The pHen6-W25 vector was transformed and a single clone was selected from the agar plates and inoculated in 20 mL of liquid LB-medium containing 100μg/ml ampicillin and 1% glucose. The bacteria were cultured at 37 °C with agitation for 16 h. The bacteria were then diluted into 1L Terrific Broth (TB) medium containing 100μg/ml ampicillin, 2mM MgCl2, 0,1% glucose and incubated at 37 °C to an OD600 of 0.6–0.9. The expression of the Nanobodies was induced by adding 1 mM of IPTG (isopropyl-β-d-1-thiogalactopyranoside) for 20h at 28°C. Bacteria were collected by centrifugation at 8,000 rpm for 8 min at 4 °C. The harvested bacteria were resuspended in a 12 mL TES buffer (0.2 M Tris pH 8.0, 0.5 mM EDTA, 0.5 M sucrose) and incubated for 1h on ice, then incubated for another hour on ice in a 18 mL TES buffer, diluted 4 times and centrifuged at 8,000 rpm at 4°C to pellet down cell debris. The supernatant was loaded on 5mL of HisPur Ni-NTA agarose resin which was pre-equilibrated with binding buffer (Tris 50mM, NaCl 500 mM, imidazole 10 mM pH 7,5). The lysed cells containing His- and myc-tagged nanobodies were added to the column followed by adding the column′s volume in binding buffer for a total of 8 times. The column was washed by adding 8-fold the column′s volume with wash buffer (Tris 50mM, NaCl 500 mM, imidazole 30 mM pH a 7.5), and eluted with 15mL of elution buffer (Tris 50mM, NaCl 150 mM, 150 mM imidazole, 1mM DTT pH 7.5). Purified Nanobodies were verified by SDS-PAGE Coomassie staining analysis.

Mammalian expression plasmids encoding SARS VHH72 and MERS VHH-55 (73) with a C-terminal a monomeric human Fc tag were transfected into ExpiCHO cells (Thermofisher) using ExpiFectamine as manufacterer’s recommendation. The supernatants were harvested on 7 days, filtered using 0.22 μM, and purified on protein A column (GE healthcare). Proteins were buffer exchanged and concentrated in PBS pH7.4.

### *In vitro* pulldown assay

100 μL of recombinant W25 (1 μg/μL) was incubated with 100 μL NHS-activated sepharose 4 Fast Flow beads coupled to SARS-CoV-2 Spike protein plus 800 μL of PBS pH 7.4 and bovine serum albumin coupled beads (negative control) for 1 h at room temperature, followed by 3 x 3 h washes with PBS-T (PBS 0.1% Tween20), 3 x 5 min washes with PBS with 500 mM NaCl and 3 x 5 min washes with PBS. The pulled down material was boiled in Laemmli sample buffer, separated by 10% SDS-polyacrylamide gels and stained with coomassie blue. Recombinant SARS-CoV-2 (2019-nCoV) Spike S1 Protein (RBD) from Trenzyme, Germany was used for the assays.

### Cell culture

HeLa cells were maintained at 37°C in DMEM/F12 supplemented with 10% FCS and 100 units/mL of penicillin and streptomycin. Plasmid transfection was performed in 10 cm plates using 25 μg of DNA, 24h after transfection cells were split into 96 well plate ∼10000 cells per well. Transfection was performed using Lipofectamine 2000 (Invitrogen) according to the manufacturer’s instructions and media were supplemented with Normocin during transfection (Invivogen).

### Dot Blot analysis screening

Individual colonies obtained from a density gradient separation protocol were inoculated into 2 mL of LB medium and incubated overnight at 37°C with 200 rpm agitation. 100 μL of pre-inoculum was added to 1.9 mL of fresh LB medium with 25 μg/mL chloramphenicol, incubated at 37°C with 200 rpm agitation until it reached OD600 of 0.45-0.6. To induce protein expression, IPTG was added at a final concentration of 50 μM for 3 h at 30°C and 200 rpm. The culture was pelleted and resuspended in 1mL PBS with 0.2% TritonX100, sonicated for 10 seconds at 40% on ice, then centrifuged at 14,000 g for 30 mins at 4°C and the supernatant was recovered to obtain a total protein extract from each clone. 1 μl of SARS-CoV2 Spike protein (200 ng/μL) and a *E. coli* total protein extract were spotted within a pre-marked grid onto a 0.2 μm pore–size nitrocellulose membrane (Merk Millipore). The membrane was left to dry to fix the proteins for 30 min at room temperature. Non-specific sites were blocked with blocking solution (PBS containing 0.1% Tween20 with 5% bovine serum albumin) for 30 min at room temperature with agitation. The blocking solution was discarded and each membrane was incubated for 1 h at room temperature with agitation at a dilution of 1:10 for the total protein extract of each clone in 5 ml of TBS-T containing 5% BSA, followed by 3 x 5 min washes with PBS-T. Secondary antibody incubation was performed with Mouse Anti-Myc antibody (9B11, Cell Signalling) (1:3000) in PBS-T containing 5% BSA for 1 h at room temperature, followed by 3 x 5 min washes with PBS-T. After this, the membrane was incubated with a Goat anti-mouse IgG antibody coupled to HRP (Invitrogen) (1:5000) in PBS-T containing 5% BSA, for 1h at room temperature, followed by 3 x 5 min washes with PBS-T and visualized using the ECL reagent (Pierce).

### High content microscopy

Spike-GFP transfected HeLa cells were grown on a 96-well optical plate (Themofisher), washed with PBS 3 times and fixed with 4% paraformaldehyde at room temperature for 30 min. After fixation, cells were washed with PBS and permeabilized in PBS 0.2% TritonX100. After washing the cells 3 times in PBS they were incubated to either an extract containing Nanobodies or purified Nanobodies during 45 minutes at 37°C. After washing another 3 times with PBS, a mouse anti-myc antibody (Cell Signaling) was used at 1:3000 and incubated during 45 minutes at 37°C. Directly afterward the cells were washed 3 times in PBS and incubated with an anti-mouse Alexa 647 during 35 minutes at 37°C. For nuclei staining, cells were washed with PBS and incubated for 10 min at room temperature with 0.1 mg/mL DAPI. After the final wash, cells were maintained in the 96-well optical plates in 100 μL PBS. Images of fixed cells were acquired with a high content automatic microscope, Celldiscoverer 7 (Carl Zeiss GmbH, Jena, Germany).

### Structural integrity and functionality tests using Tycho NT.6

The binding of W25 and the structural integrity of Spike RBD was verified using a label-free thermal shift assay with Tycho NT.6 (Nanotemper Technologies GmbH) using intrinsic tryptophan and tyrosine fluorescence. 10 μL solutions of Spike RBD (2 μM), W25 (2 μM) and Spike RBD mixed with W25 (2 μM each) were prepared and loaded into Capillaries Tycho NT.6 (TY-C001, NanoTemper Technologies GmbH). The Tycho instrument applied a quick thermal ramp from 35°C to 95°C with a heating rate of 30°C/min and the unfolding of proteins were monitored through changes in the 350 nm / 330 nm fluorescence emission ratio. Recombinant SARS-CoV-2 (2019-nCoV) Spike S1 Protein (RBD) from Trenzyme, Germany was used for the assays.

### Binding affinity measurements using MST

The dissociation constant (Kd) between the W25 Nanobody and SARS-CoV-2 S1 of Spike RBD was measured by microscale thermophoresis (MST) using a Monolith NT.115PICO instrument (Nanotemper Technologies GmbH). Purified W25 was buffer exchanged into a PBS buffer, pH 7.4, and its concentration was adjusted to 10 μM using UV-absorbance. Next, W25 was fluorescently labeled with the Monolith Protein Labeling Kit RED – NHS 2nd Generation (MO-L011, NanoTemper Technologies GmbH) following the protocol established in the manual. Labeled W25 was centrifuged at 14,000 rpm for 15 min to eliminate precipitates. A 16-point serial dilution series of recombinant Spike RBD (250 nM – 7.6 pM) was applied in PBS buffer containing 0.01% Pluronic F-127 and mixed with a final concentration of 1 nM labeled W25. Affinity measurements were conducted in Premium Capillaries Monolith NT.115 (MO-K025, NanoTemper Technologies GmbH) and repeated three times. Recombinant SARS-CoV-2 (2019-nCoV) Spike S1 Protein (RBD) from Trenzyme, Germany was used for the assays.

### Plaque Reduction Neutralisation (PRNT) assay

SARS-CoV-2 isolate QLD02/2020 – 30/1/2020 (GISAID accession EPI_ISL_407896) and QLDID935/2020 – 25/03/2020 (GISAID accession EPI_ISL_436097), referred as D614 and G614, respectively, was isolated and obtained from Queensland Health, Brisbane, Australia. Viruses were passaged three times in Vero E6 cells and titrated by focus-forming assay on Vero E6 cells. Serially dilutions of purified nanobody or nanobody fused monomeric Fc were mixed with ∼250 FFU/well of SARS-CoV-2 viruses and incubated for 1 h at 37°C. Subsequently, mixtures were added to previously-plated E6 monolayer cells and incubated at 37°C for 30 min. Cells were then overlaid with 1% (w/v) medium viscosity carboxymethyl cellulose in M199 (Gibco) supplemented with 2% heat-inactivated fetal bovine serum (Bovogen) supplemented with 1% Penicillin-Streptomycin (Gibco) and incubated at 37oC in 5% CO2. After 14 hrs incubation, overlay was removed, and cells fixed with 80% cold-acetone in PBS for 30 min at -20°C. Plates were then dried, blocked with blocking buffer containing 1xKPL (Seracare) and 0.1% PBS-Tween 20 for 1 h and then incubated with 1 μg/ml of human CR3022 anti-spike mAb and followed by 0.2 μg/ml IR-Dye800-conjugated goat anti-human IgG (Millienium Science) in blocking buffer. Plates were washed 3 times after antibody incubations by submerging in PBS-T 0.1%Tween-20. Plates were then dried prior to visualizing using Odyssey (LI-COR). Immunoplaques were manually counted. The neutralizing antibody titers were defined as the amount of antibody (nM) resulting in a 50% reduction relative to the total number of plaques counted without antibody.

## Author Contributions Statement

Immunization and animal wellbeing T.P, B.U and C.D; Bacterial display and gradient protocol development G.V.N, R.J, J.H, C.S, T.P, N.L.G.R, Z.M.C, Y.C, A.C, L.A.F, Y.M, S.G.M, H.M, A.M, B.U, P.K, P.E, J.B, P.C.C, L.A.F & A.R.F; SARS-CoV-2 PBMCs isolation T.P & J.H; libraries construction J.H & R.J; Nanobodies selection Y.M, L.A.F, R.J & G.V.N; Nanobodies binding confirmation G.V, J.H & A.R.F; Nanobodies expression Y.C, R,J, D.W, N.M, A.A.A; Neutralization and live virus work D.W, N.M, A.A.O & A.A.K; Conceptualization D.B, A.B, A.R.F, G.V, R.J, D.S, L.A.F & C.S; Data curation, D.B, N.M, A.A.O, A.B, G.V.N, R.J, J.H, C.S, T.P, N.L.G.R, Z.M.C, Y.C, A.C, A.R.F; Analysis A.B, A.R.F, G.V & R.J; Funding acquisition D.W, P.K, J.B, A.M, B.U, P.C.C, G.R & A.R.F; Resources D.W, J.B, A.M, B.U, P.C.C, D.S, G.R & A.R.F; Supervision A.R.F; Writing—original draft, D.W, A.B, L.A.F, A.R.F, D.S, G.V & R.J; Writing—review and editing, D.S, D.W, N.M, A.A.O, G.V.N, R.J, J.H, C.S, T.P, N.L.G.R, Z.M.C, Y.C, A.C, L.A.F, A.B, Y.M, S.G.M, A.M, B.U, P.K, P.E, G.R, A.A.K & A.R.F. All authors have read and agreed to the published version of the manuscript.

### Acknowledgment

The authors would like to thank the KOICID for partially supporting this study. We thank Dr. Ronald T. Hay for his generous support and assistance. This work was funded by FONDECYT No. 11150532 to ARF; FONDEF No ID17I10037 to ARF; FONIS EU-LAC T010047 to PCC, JO, JB, NLR & ARF; PAI-CONICYT No 79150075 to ARF; FONDEQUIO EQM180037 to ARF; the regional Council of the “Los Rios region” project FICR16-19 and FICR19-20; J.B was funded by ISCIII Miguel Servet Program CP19/00200. Graduate fellowship ANID N° 21161365 to R.J; ANID N°21160481 to A.C; ANID N° 22170632 to C.S; Z.M.C was funded by the academic graduate students fellowship of the University of Costa Rica of the academic mobility program; N.L.G.R was funded by Becas Santander Iberoamerica Investigacion 2018/2019; H.M was funded by a postdoctoral fellowship from FONDECYT 3170159. Work in the laboratory of L.A.F is supported by Grants BIO2017-89081-R (Agencia Española de Investigación AEI/MICIU/FEDER, EU) and CSIC PIE-RDL-COVID-19 (Ministerio de Ciencia e Innovación from Spain). We thank Dzongsar Khyentse Rinponche for his generous donation of alpacas to this project. We would like to thank Amber Philp for proofreading and Rocio Miranda, Andrea Bastidas, Vanessa Wiederhold, and Viviana Estrada for administrative support, Felipe Serrano for illustration and Omer Navarrete for his commitment to the wellbeing of the alpacas.

