## Supplementary figures and images for "Potent neutralization of clinical isolates of SARS-CoV-2 D614 and G614 variants by a monomeric, sub-nanomolar affinity Nanobody"

### Supplemental Figure-1

#### Supplemental Figure- 1

a

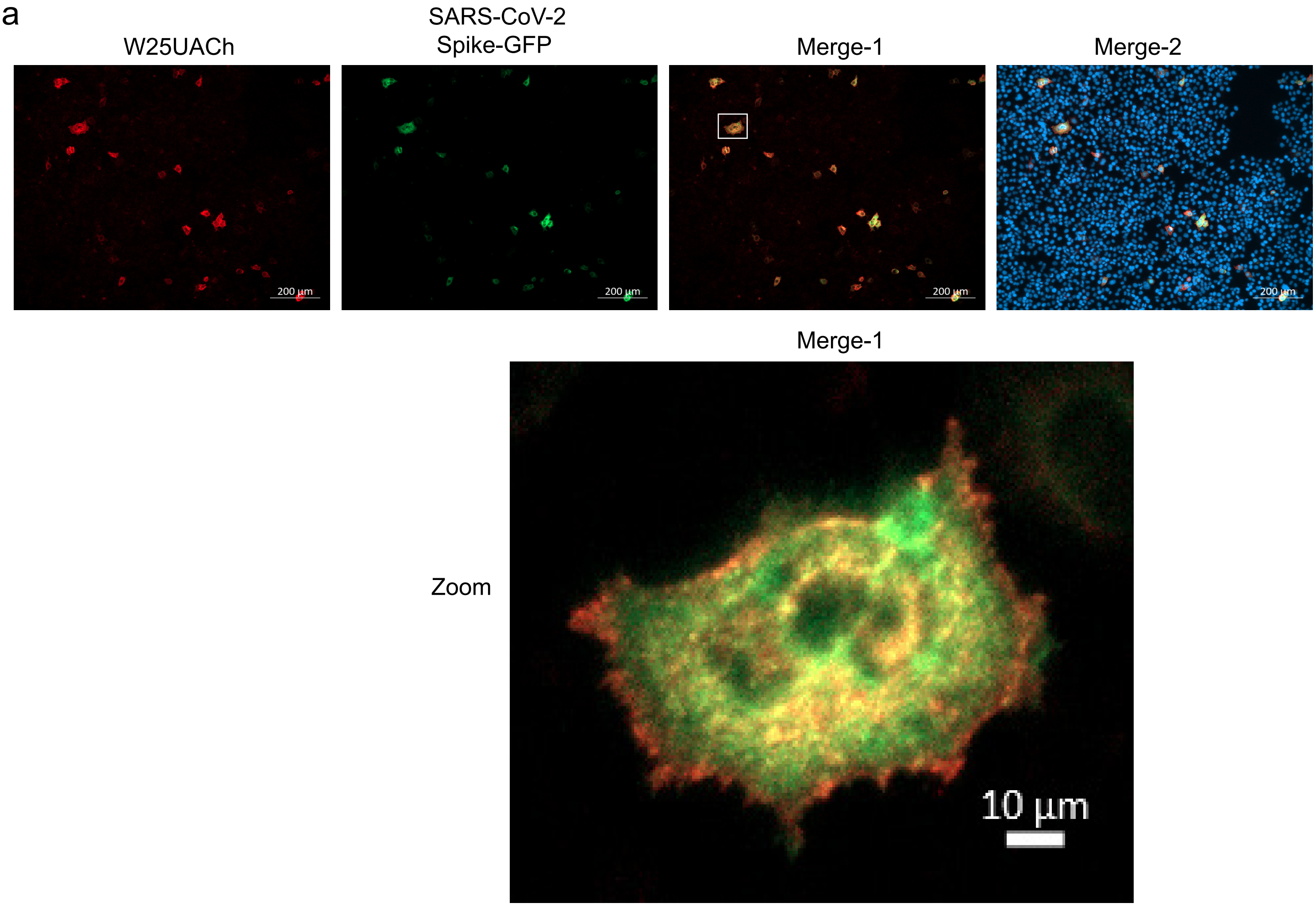


b

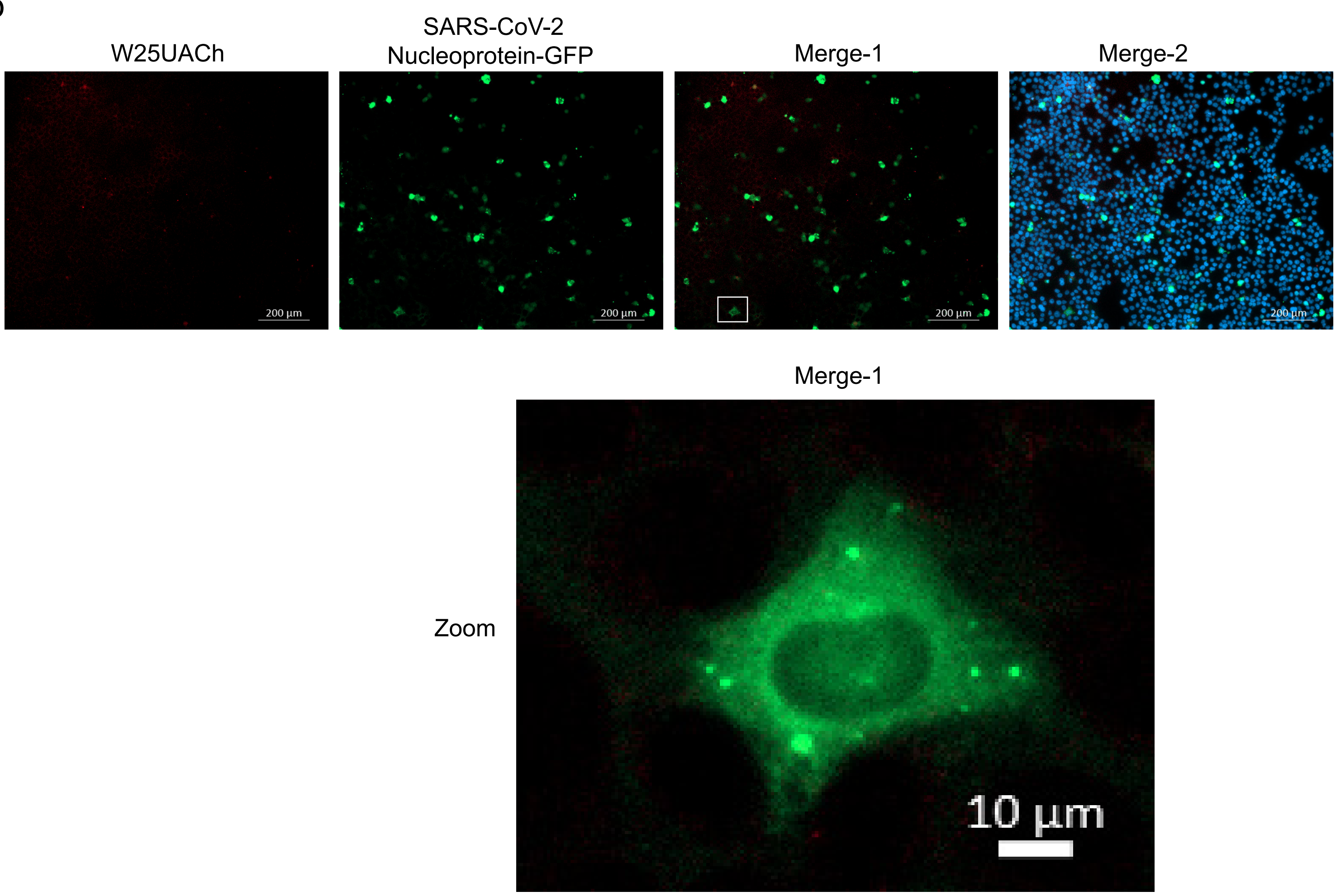
