## Supplemental information for "Potent neutralization of clinical isolates of SARS-CoV-2 D614 and G614 variants by a monomeric, sub-nanomolar affinity Nanobody"

**Supplemental Figure Legends**

**Supplemental Figure-1. Controls and expression**. **A)** Immunofluorescence assays of Spike-GFP transfected HeLa cells using purified recombinant W25UACh nanobody as primary antibody. **B)** Immunofluorescence assays of Nucleoprotein- GFP transfected HeLa cells using purified recombinant W25UACh nanobody as primary antibody**.**

### Nanobodies Sequences

#### Nucleotide sequences W23 Nanobody

ATGGCTCAGGTGCAGCTGGTGGAGTCTGGGGGAGGCTTGGTGCAGCCTGGGGAGTCTCTGAGACTCTC
CTGTGCAGCCTCTGGAAACATCTTCGGAATCGCTGCCGTGCACTGGTTCCGCAAGGCTCCAGGGAAGGA
GCGCGAGTTTACTGCAGGTTTTGGTAGTGATGGTAGCACAAACTATGCAAACTCCGTGAAGGGCCGATT
CACCATCTCCAGAGACAATGCCAAGAACACGACATATCTGCAAATGAACAGCCTGAAACCTGAGGACAC
GGCCGTCTATTATTGTCATGCGCTAATCAAGAATGAACTTGGATTCCTTGATTACTGGGGCCCGGGGACC
CAGGTCACCGTCTCCTCA

#### Amino acid sequences W23 Nanobody

MAQVQLVESGGGLVQPGESLRLSCAASGNIFGIAAVHWFRKAPGKEREFTAGFGSDGSTNYANSVKGRFTIS
RDNAKNTTYLQMNSLKPEDTAVYYCHALIKNELGFLDYWGPGTQVTVSS

#### Nucleotide sequences W25 Nanobody

ATGGCTCAGGTGCAGCTGGTGGAGTCTGGGGGAGGCTTGGTGCAGCCTGGGGAGTCTCTGAGACTCTC
CTGTGCAGCCTCTGGAAGTATCTTCGGAATCTATGCCGTGCACTGGTTCCGCATGGCTCCAGGGAAGGA
GCGCGAGTTTACTGCAGGTTTTGGAAGTCATGGTAGCACAAATTATGCAGCTTCCGTGAAGGGACGATT
CACCATGTCCAGAGACAATGCCAAGAACACGACGTATCTGCAAATGAACAGCCTGAAACCTGCGGACAC
GGCCGTCTATTACTGTCATGCGCTAATAAAGAATGAACTTGGGTTCCTTGACTACTGGGGCCCGGGGAC
CCAGGTCACCGTCTCCTCA

#### Amino acid sequences W25 Nanobody

MAQVQLVESGGGLVQPGESLRLSCAASGSIFGIYAVHWFRMAPGKEREFTAGFGSHGSTNYAASVKGRFT
MSRDNAKNTTYLQMNSLKPADTAVYYCHALIKNELGFLDYWGPGTQVTVSS
